# ^15^N metabolic labeling quantification workflow in Arabidopsis using Protein Prospector

**DOI:** 10.1101/2021.11.30.470624

**Authors:** Ruben Shrestha, Andres V. Reyes, Peter R. Baker, Zhi-Yong Wang, Robert J. Chalkley, Shou-Ling Xu

## Abstract

Metabolic labeling using stable isotopes is widely used for the relative quantification of proteins in proteomic studies. In plants, metabolic labeling using ^15^N has great potential, but the associated complexity of data analysis has limited its usage. Here, we present the ^15^N stable-isotope labeled protein quantification workflow utilizing open-access web-based software Protein Prospector. Further, we discuss several important features of ^15^N labeling required to make reliable and precise protein quantification. These features include ratio adjustment based on labeling efficiency, median and interquartile range for protein ratios, isotope cluster pattern matching to flag incorrect monoisotopic peak assignment, and caching of quantification results for fast retrieval.

## 1. Introduction

Accurate and high-throughput protein quantification is fundamental to proteomic studies (Aebersold and Mann, 2003). To provide the highest quantification accuracy when comparing samples one needs to minimize differences introduced in the processing of samples and acquiring the data. This can be best achieved through the introduction of stable isotopes into samples that allow samples to be mixed and then analyzed in the mass spectrometer. The application of metabolic labeling, which uses stable-isotope labeled amino acids in cell culture (SILAC)(Ong et al., 2002) or ^15^N nitrogen-containing salts (Guo and Li, 2011; Schulze and Usadel, 2010) into the whole cell or organism *in vivo*, enables relative quantifications of proteins on a global scale. In such a quantitative experiment, one sample is labeled with the natural abundance (light), and the other with a stable isotope of low natural abundance (heavy) during growth. The samples are mixed, processed, and analyzed by the mass spectrometer. Chemically identical peptides from these light- and heavy-labeled mixed samples co-elute by chromatography into the mass spectrometer, which can distinguish between the light and heavy peptides based on their mass difference, and thereby quantify the difference in peptide, and hence protein abundance between the samples. An alternative stable isotope-based strategy is to chemically tag peptides after enzymatic digestion; the most popular reagents for this strategy are isobaric tagging reagents Tandem Mass Tags (TMT) (Pappireddi et al., 2019). The TMT isobaric tagging reagents allow comparison of a larger number of samples, but the labeling is done at the peptide level after sample digestion and then samples are mixed. In contrast, the metabolic labeling is introduced into the proteins during growth, thus samples can be combined at the beginning, minimizing variations introduced by sample processing that can compromise quantification accuracy (Piehowski et al., 2013).

Although SILAC has been widely used in animal cell lines and has been the gold standard for MS-based proteomics quantification (Schubert et al., 2017), ^15^N-labeling based quantitative applications are still quite limited in plants despite it being cheaper (Arsova et al., 2012). This could be due to the complexity of the data analysis. SILAC pairs are easily identifiable because they have well-defined mass differences as typically only lysine and arginine are labeled. In contrast, in ^15^N labeling, each amino acid in the expressed proteins is labeled, and therefore, the mass difference in ^15^N pairs varies depending on the number of nitrogen atoms in their composition. Also, as more amino acids are being labeled, the effect of incomplete incorporation of the heavy isotope can be more pronounced under some conditions, such that isotope clusters of heavy labeled peptides in the survey scan MS^1^ spectra are generally broader, making it harder to identify the monoisotopic peak. This can lead to significantly reduced identification of heavy labeled peptides.

There are very few freely available software tools with workflows that can analyze large-scale ^15^N labeled samples. Such tools include MSQuant (Mortensen et al., 2010), pFIND (Li et al., 2005), and Protein Prospector (Chalkley et al., 2005; Li et al., 2005). The workflow using MSQuant normally requires manual inspection of the pairs of the light and heavy forms that both fit with expected isotope envelope distribution; those that don’t fit the criteria will be omitted from further analysis (Kierszniowska et al., 2009). This makes it very time-consuming for a large dataset because of the manual inspection prerequisite. In addition, if both forms need to be present for quantification, then there will be a high false-negative rate for some of those highly biologically interesting proteins which only express in one of the two conditions, or from immunoprecipitated (IP) samples where those proteins will be only in the bait-IP but may be completely absent in the control IP.

Here, we present the ^15^N quantification workflow based on the free web-based software Protein Prospector (Chalkley et al., 2005; Li et al., 2005). After data search with respective ^14^N and ^15^N search parameters, quantification between the light and heavy peptide pairs is done based on the identification of either the light or heavy peptide, or both. The calculated peptide ratio is then adjusted based on the labeling efficiency input. Peptide ratios are then compiled into protein-level statistics such as median and interquartile ranges. Additional features in Protein Prospector include a Cosine Similarity (CS) score which can be utilized to reduce manual checking of spectra and a cache function that enables efficient result retrieval through cached result storage. This workflow allows us to report quantifications of thousands of proteins and is applicable to the quantification of the total proteomes, sub-proteomes, and immunoprecipitated samples (Bi et al., 2021; Garcia et al., 2020; Park et al., 2019). It can be also applied to the quantification of post-translational modification with a slight modification.

## 2. Materials and Equipment

The materials and data acquisition for the first Col/*acinus-2 pinin-1* dataset were detailed in (Bi et al., 2021). Briefly, the wild type (Col) and *acinus-2 pinin-1* plants were grown on Hoagland medium containing ^14^N or ^15^N (1.34 g/L Hogland’s No. 2 salt mixture without nitrogen, 6 g/L Phytoblend, and 1 g/L KNO_3_ or 1 g/L K^15^NO_3_ (Cambridge Isotope Laboratories), pH 5.8) for 14 days under the constant light condition at 21–22 °C on vertical plates. Proteins were extracted from six samples (one ^14^N-labeled Col, two of ^15^N-labeled Col, two of ^14^N-labeled *acinus-2 pinin-1*, and one ^15^N-labeled *acinus-2 pinin-1*) individually using SDS sample buffer and mixed as the following: one forward sample F1 (^14^N Col/^15^N *acinus-2 pinin-1*) and two reverse samples R1 and R2 (^14^N *acinus-2 pinin-1*/^15^N Col) and separated by the SDS-PAGE gel with a very short run (∼3 cm). Two segments (upper part (U) ranging from the loading well to ∼50 KD; lower part (L) ranging from ∼50 KD to the dye front) were excised, trypsin digested, and analyzed by liquid chromatography-mass spectrometry (LC-MS) as described in (Bi et al., 2021) on a Q-Exactive HF instrument using 50 cm column ES803.

For the second Col/*sec-5* dataset, the WT and *sec-5* plants were grown on Hoagland medium containing ^14^N or ^15^N (1.34 g/L Hogland’s No. 2 salt mixture without nitrogen, 6 g/L Phytoblend, and 1 g/L KNO_3_ or 1 g/L K^15^NO_3_ (Cambridge Isotope Laboratories), pH 5.8, 1% Sucrose) for 14 days. Plates were placed vertically in a growth chamber under the constant light condition at 21– 22 °C. Whole plant tissues were harvested in liquid nitrogen. Proteins were extracted from eight samples (two ^14^N-labeled Col samples - 1 and 5; two of ^15^N-labeled Col samples - 2 and 6; two of ^14^N-labeled *sec-5* samples - 3 and 7; and two ^15^N-labeled *sec-5* samples - 4 and 8) individually using 2X SDS sample buffer (plant tissue mass (mg): buffer (µL) ratio=1:3) and mixed as the following: two forward samples F1 and F2 (^14^N Col/^15^N *sec-5*, Mix1+4; Mix 5+8) and two reverse samples R1 and R2 (^14^N *sec-5*/^15^N Col, Mix 2+3; Mix 6+7) and separated by the SDS-PAGE. Five segments were excised, and trypsin digested. The peptide mixtures were desalted using C18 ZipTips (Millipore) and analyzed by liquid chromatography-mass spectrometry (LC-MS) the same as above described in (Bi et al., 2021) on a Q-Exactive HF instrument using 50 cm column ES803.

The following items are required for data analysis in this method.

1. A personal computer with internet access and 8 GB or above of RAM.
2. A web browser.
3. Protein Prospector (https://prospector.ucsf.edu/prospector/mshome.htm). We recommend using a local installation for quantification as it needs to access the raw data. A version for local installation is provided for free upon request through the above website.
4. The Thermo Raw FileReader package.
5. Peak list generation software (e.g. MSConvert, part of the Proteowizard package that can be downloaded for free)
6. Image J (can be download for free from https://imagej.nih.gov)

## 3. Method

### 3.1 Overview of the Procedure

This section outlines the major steps of ^15^N Identification and quantification (ID & Quan) analysis. A detailed step-by-step protocol is provided in the procedure below. The ID & Quan analysis involves four major steps as listed in Figure 1 workflow.

**Figure 1:**
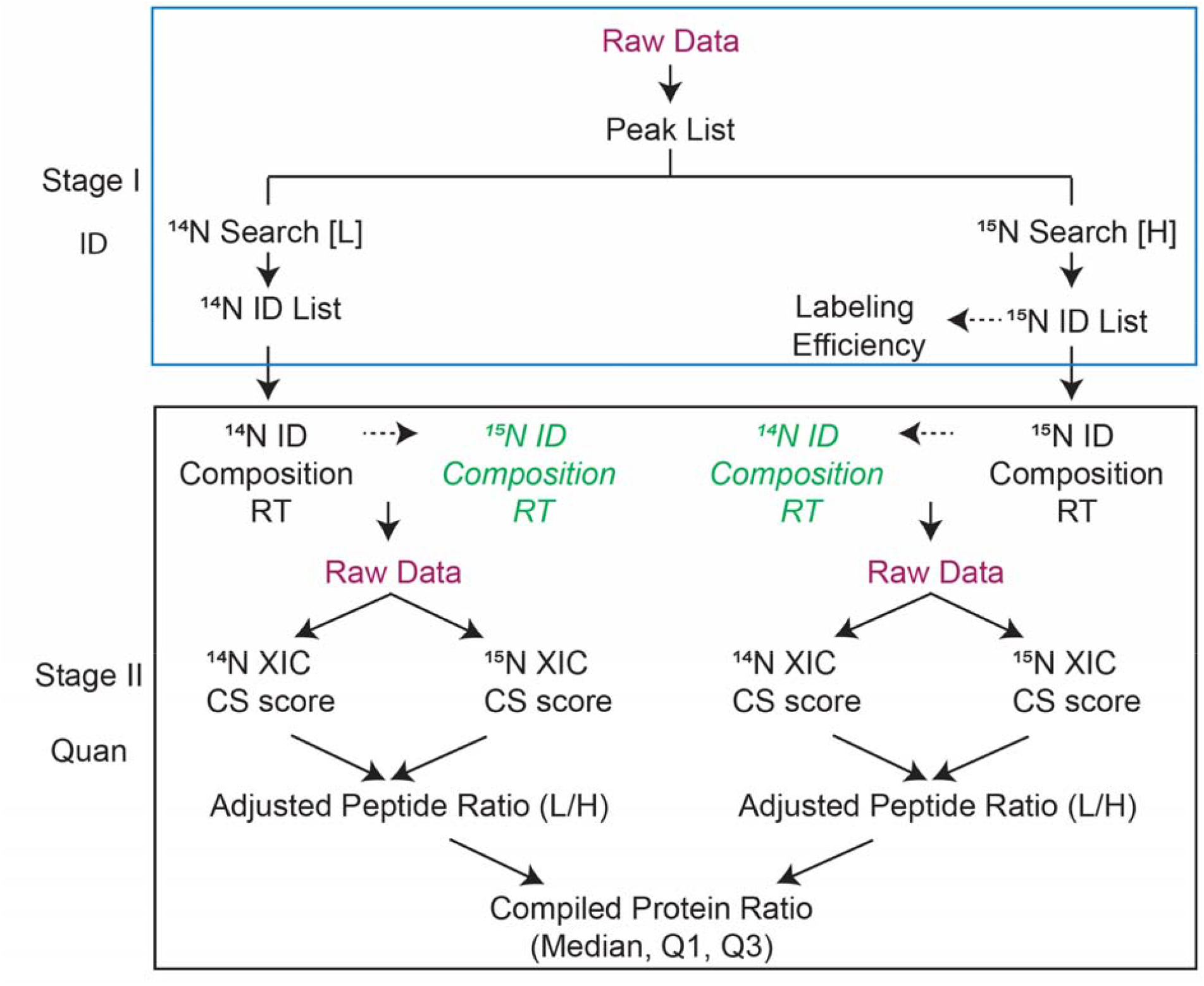
Workflow of ^15^N metabolic labeling protein quantification used in Protein Prospector. There are two stages: Stage I-Identification of proteins; Stage II-Quantification of proteins/peptides. Labeling efficiency is determined by the user and then input as a parameter to correct ratios to generate adjusted peptide ratios. Based on the identification of peptides either from ^14^N search [L] or ^15^N search [H], the PP software finds the matching counterpart and retrieves isotope cluster intensities and/or areas. The ratios between the peptide pair are reported and compiled into protein-level statistics such as median and interquartile ranges. CS scores are calculated on the matched pairs based on the comparison of experimental vs theoretical isotope distributions.

1. Identify ^14^N and ^15^N labeled proteins by searching the data with corresponding parameters separately.
2. Determine the labeling efficiency or enrichment of ^15^N labeled peptides by comparing the experimental to the theoretical peak isotope profile for peptides with different labeling efficiency.
3. Submit quantification in “*Search Compare”* with user-specified parameters to extract quantification information.
4. Retrieve the report for the quantification and (optional) determine the quality of quantification and follow up with informatics analysis.

### 3.2 Step 1: Identification: Search and Identify ^14^N and ^15^N labeled proteins

The peak lists are first generated from the raw files using in-house peaklist generator script PAVA (Guan et al., 2011a) and deposited into the folder, which is a mirror of the raw file folder, and searched against the Arabidopsis database. A project name is created for the search. Files from the same experiment can be searched together under the same project name with separate search parameters for ^14^N and ^15^N peptides as listed in table 1.

**Table 1:** Search parameters for ^14^N light- and ^15^N heavy-labeled searches. The following parameters apply to data acquired on high-resolution Orbitrap data for both MS^1^ survey scan and MS^2^ fragment scan and can be easily adjusted to data acquired on other instruments if necessary.

| Parameter Name | Setting |
| --- | --- |
| Database | *TAIR10_pep_20101214.fasta.random.concat* |
| Taxonomy | All |
| Masses are | monoisotopic |
| Parent (MS1) Tolerance | 10 ppm |
| Frag (MS2) Tolerance | 20 ppm |
| Digest | Trypsin |
| Max. Missed Cleavages | 1 |
| Max. Mods | 2 |
| **N14 specific search parameters ($^{14}\text{N}$ search [L]):** |  |
| Constant Modification | Carbamidomethyl (C) |
| Variable Modification | Acetyl (Protein N-term); Acetyl+Oxidation (Protein N-term M);<br>Gln → pyro-Glu (N-term Q); Met-loss (protein N-term M);<br>Met-loss+Acetyl (Protein N-term M); Oxidation (M). |
| **N15 specific search parameters ($^{15}\text{N}$ search [H]):** |  |
| Constant Modification | Label: 15N(1) (A); Label: 15N(1) (D);<br>Label: 15N(1) (E); Label: 15N(1) (F); Label: 15N(1) (G);<br>Label: 15N(1) (I); Label: 15N(1) (L); Label: 15N(1) (M);<br>Label: 15N(1) (P); Label: 15N(1) (S); Label: 15N(1) (T);<br>Label: 15N(1) (V); Label: 15N(1) (Y);<br>Label: 15N(1)+Carbamidomethyl(C);<br>Label: 15N(2) (K); Label: 15N(2) (N); Label: 15N(2) (Q);<br>Label: 15N(2) (W); Label: 15N(3) (H); Label: 15N(4) (R) |
| Variable Modification | Acetyl (Protein N-term); Met-loss (protein N-term M);<br>Met-loss+Acetyl (Protein N-term M);<br>Label: 15N(1)+Acetyl+Oxidation (Protein N-term M);<br>Label: 15N(1)+Gln → pyro-Glu (N-term Q);<br>Label: 15N(1)+Oxidation (M). |

**Note:**
- 1) If the samples are enriched for post-translational modifications (PTMs), users need to include these in the variable parameters for both the $^{14}\text{N}$ search [L] and $^{15}\text{N}$ search [H] search. - 2) We recommend both the MS<sup>1</sup> survey and MS<sup>2</sup> fragment scans be acquired with high resolution.

### 3.3 Step 2: Determine the labeling efficiency or enrichment

The ^15^N labeling can often be incomplete, but typically constant across all proteins in a given experiment (Skirycz et al., 2011). However, the labeling efficiency between different experiments can range between 93-99% after 14 days of labeling on plates or in liquid culture for Arabidopsis plants, depending on the chemical used, labeling duration (the number of the plant cell doubling), and the availability of the nitrogen. If a sample has 95% labeling enrichment, it means the ^15^N labeled peptide has 95% of ^15^N and 5% standard ^14^N.

Because Protein Prospector uses only the monoisotopic peak (M) for quantification, the accuracy of quantification will be affected if the ratio is not corrected for the labeling efficiency. To do this, PP allows estimation of the labeling efficiency in the “MS-Isotope” module. MS-Isotope allows plotting of the theoretical isotope distribution pattern for user-defined levels of heavy isotope incorporation (Supplemental Figure 1), and the user will manually compare these plots to observed distributions to determine the labeling efficiency.

Figure 2 and supplemental Figure 1 and 2 show how labeling efficiency is determined in one experiment. Figure 2A shows theoretical pattern computed for a range (95 to 99%) of labeling efficiency, and notably, M-1/M ratio is most affected by different labeling efficiency (Schaff et al., 2008), and thus this ratio is used to calculate the actual incorporation rate. Peptides with small mass (m <1500) are more preferred for calculation of incorporation, because the M peak of such peptides is the largest peak and can be set as 100% maximum for easy calculation, and M-1/M peak is reversely correlated with the labeling efficiency (Figure 2B) (Schaff et al., 2008).

**Figure 2:**
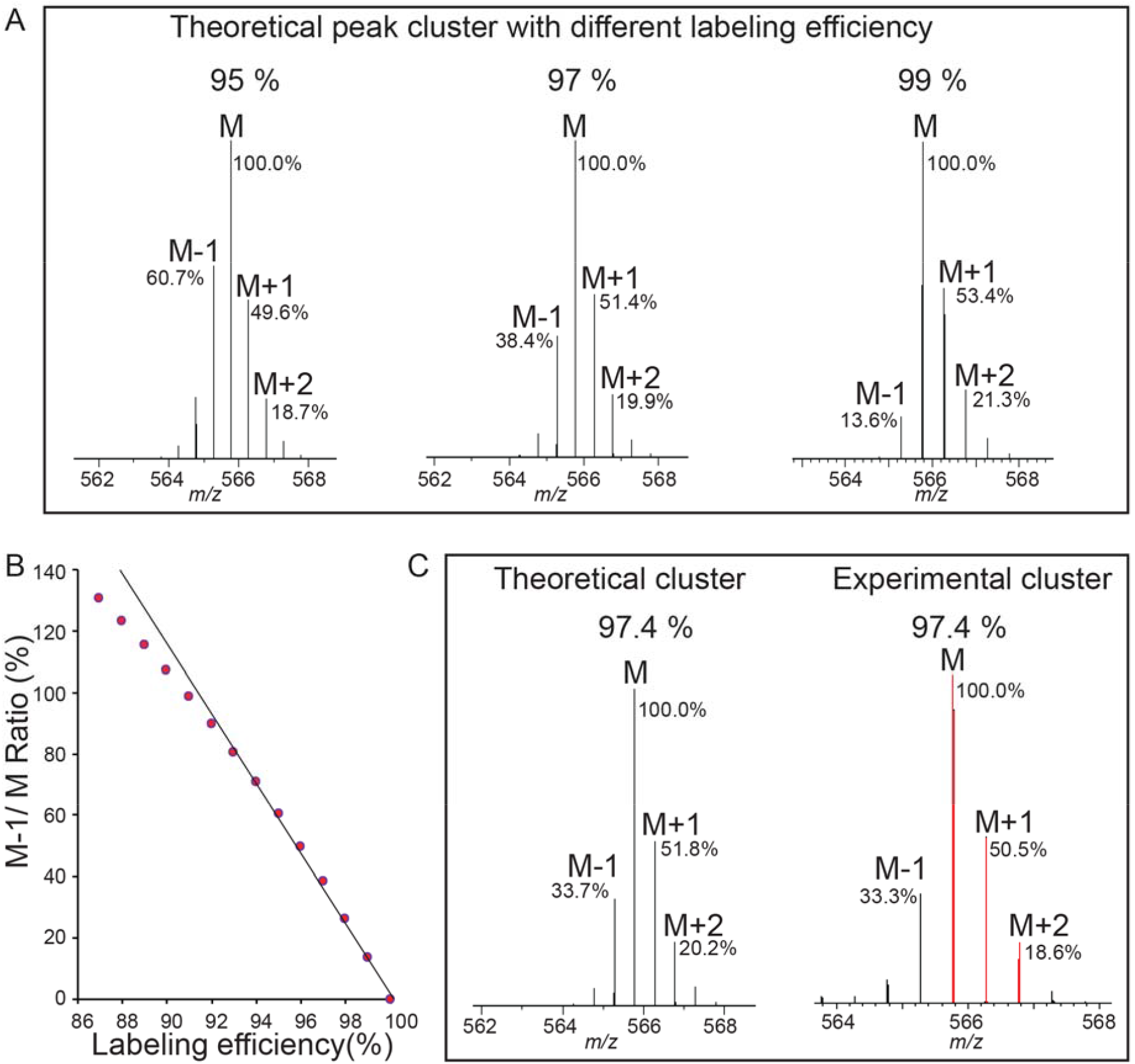
Labeling efficiency or enrichment is determined on ^15^N labeled peptides by comparing the experimental isotope pattern to the theoretical isotope profile. (A) Three plots show the theoretical isotope profile of the heavy peptide “VALEAC*VQAR” (C46 H82 O15 15N14 N1 S1) from RIBULOSE-BISPHOSPHATE CARBOXYLASES labeled at 95%, 97%, and 99% labeling efficiency (C*,Carbamidomethylation). The monoisotopic peak is the most intense peak in the isotope cluster for this *m/z* and any peaks to its left are caused by incomplete labeling. The relative abundances of M-1, M, M+1, M+2 to the M peak are labeled, with more abundance of M-1 peak indicating lower labeling efficiency. The analysis is performed using the Protein Prospector “MS-Isotope” module. (B) M-1/M ratio is reverse correlated with labeling efficiency using same peptides in (A) for calculation. (C) Labeling efficiency is determined as 97.4%.

To calculate M-1/M ratio, one easy way is to copy the image of the experimental peaks from Protein Prospector to imageJ. The intensity of monoisotopic peak (M) is set as known distance of 1, then M-1 peak intensity is measured and the percentage ratio of M-1/M is calculated, which is used to compare to theoretical numbers using MS-Isotope (Supplemental Figure 1) so to determine the labeling efficiency. Multiple peptides from different abundant proteins should be examined. We found in general examining about 8-10 peptides from different proteins (Supplemental Figure 2) is sufficient because the labeling efficiency within an experiment is relatively constant (Schaff et al., 2008). Based on these data, the overall labeling efficiency for this experiment is averaged around 97.0% (Supplemental Figure 2). Such manual calculation could be automated by the PP in the future.

By providing this labeling efficiency as a parameter when calculating the quantification, PP determines what percentage of the isotope cluster for a given peptide should be present in the monoisotopic peak, and therefore, the abundance of the satellite peaks (before M peaks due to incomplete labeling) is added to the total peptide abundance for ratio calculation.

### 3.4 Step 3: Perform quantification

Identification of at least one version of peptide (^14^N or ^15^N) in the pair of the peptides is necessary for quantification. For instance, if the peptide sequence is identified in the ^14^N search, Protein Prospector will compute the heavy-labeled peptide’s expected sequence and composition and derive the mass and isotope distribution. To improve the quality of the quantification, Protein Prospector can average together scans over a time window around when the peptide was identified to improve the signal to noise. It can use peak intensities and/or peak areas for ratio calculation based on the monoisotopic peak. The ratio reported by Protein Prospector is in the format of light divided by heavy (L/H) no matter whether the peptide is identified from ^14^N search or ^15^N search.

Figure 3 displays example Search Compare parameters for quantification. The parameters can vary depending on the experimental goals and are user-specifiable.

**Figure 3:**
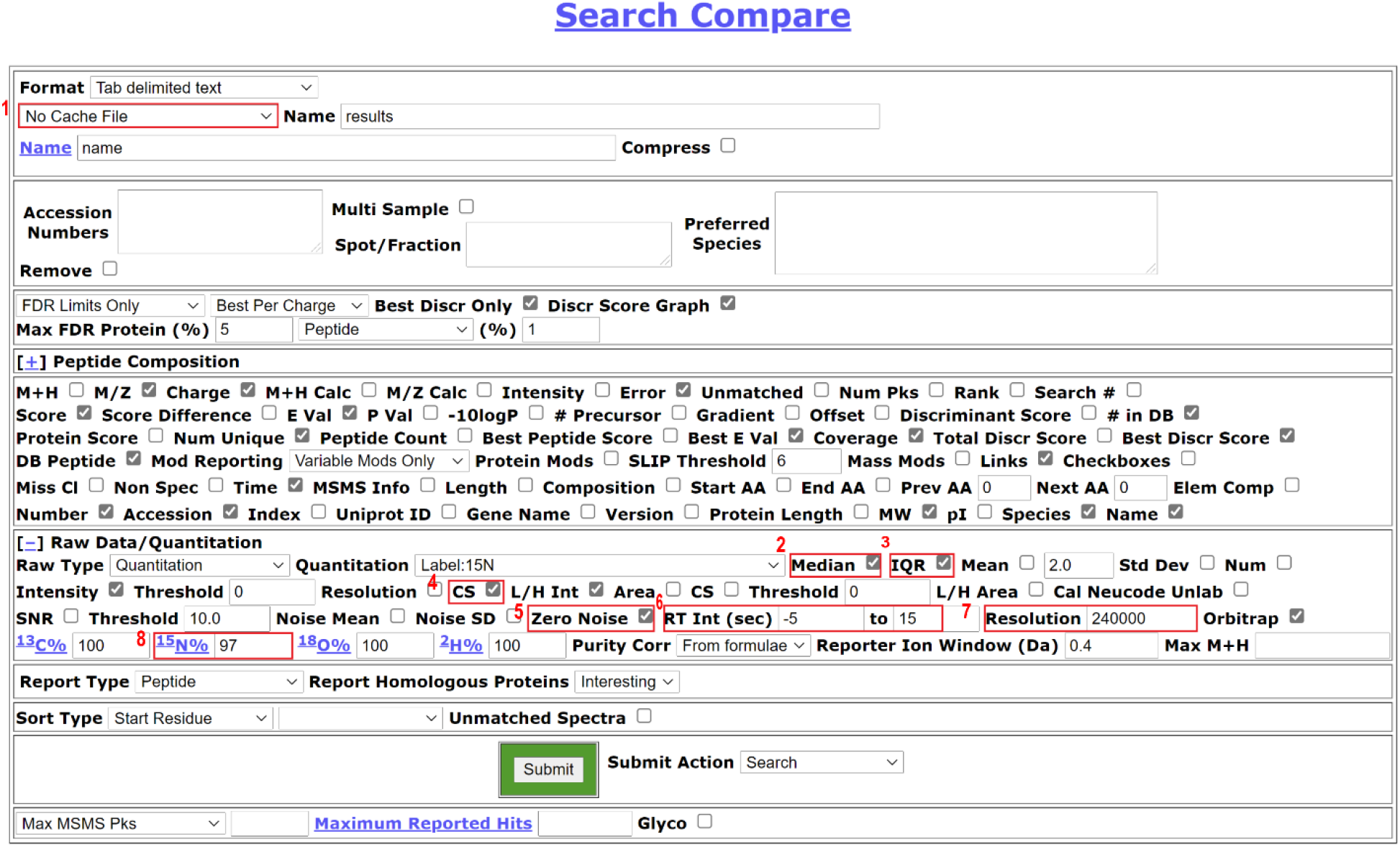
Search Compare parameters suitable for ^15^N quantification. The users can generate different formats of reports depending on the next step analysis. Users can choose a cache function in the drop-down, as highlighted in [1], which allows the users to store the analysis result and retrieve the analysis data quickly in the future. For protein quantification, we recommend checking median [2] and IQR [3] to report a median ratio of peptides for a protein and the interquartile range. CS score [4] is selected to check the quality of the match to the isotope cluster. Zero noise [5] should be checked for high-resolution MS^1^ data so the PP will not impute numbers based on the local noise level. Retention (RT) interval between −5 to 15s [6] is the window before and after when the MS^2^ scan was acquired to use for averaging MS1 spectra before calculating peak ratios. The intensity of the respective peptides is averaged across the window and the L/H ratio is reported. Resolution [7] and labeling efficiency [8] are manually filled based on the resolution setting for the MS^1^ data in the instrument settings and the data labeling efficiency decided in the previous step of this workflow, respectively.

Note:

1. In our data, peptide chromatographic peak widths are about 30 seconds, but using a narrow window (−5 to +15s) in general gives better quantification results as this nearly always includes the apex of elution but is less prone to co-eluting peptide interference. The peak width is dependent on the column and the gradient, and should be empirically examined.
2. While our data is typically acquired at 120,000K resolution, we find a 240,000K resolution setting in PP gives us more consistent quantification data to results using *Skyline* for quantification (Schilling et al., 2012) (parameters: ± 5 ppm cut-off in centroid mode).

## 4. Results and Discussion

### 4.1 Large-scale data analysis and quantification

This workflow can quantify thousands of proteins simultaneously. We demonstrate its performance using three datasets listed in the table described in (Bi et al., 2021). Previously, we had shown ACINUS and PININ genes regulate transcription and alternative splicing and we hypothesized some proteins have altered expression in the double mutant. Therefore, MS experiments were designed to identify these altered proteins on a global scale.

Only two organisms can have very good labeling efficiency within a short period (1-2 weeks): algae (Kim, unpublished) and Arabidopsis (Garcia et al., 2020), have so far been shown to have a good labeling efficiency with 98-99% efficiency being possible. In contrast, tomatoes can achieve 99% labeling efficiency after 2 months of growth (Schaff et al., 2008). However, while we consistently observe 98-99% labeling efficiency in algae (Kim, unpublished), we have observed that labeling in Arabidopsis plants is more variable, ranging from 93-99% depending on the experiment. Within one experiment, the labeling efficiency in different proteins is relatively constant (supplemental Figure 2). With less complete labeling, the identification rate of heavy labeled peptides is significantly lower than light due to errors in monoisotopic peak assignment (Table 2). If the labeling efficiency is achieved 98.5% or above, the identification rate between ^14^N and ^15^N search is similar in our experience.

**Table 2:** Summary of the identification and quantification data from seven biological experiments. The median ratio of top 100 proteins quantified (L/H) in each experiment is calculated for normalization as samples are not mixed exactly at 1:1 ratio.

| Experiments | $^{14}\text{N}$ identified proteins | $^{15}\text{N}$ identified proteins | Labeling efficiency | Quantified proteins ($^{14}\text{N}$ and $^{15}\text{N}$ combined) | The median ratio of top 100 proteins quantified (L/H) |
| --- | --- | --- | --- | --- | --- |
| Forward labeling (F1)<br>($^{14}\text{N}$ Col/$^{15}\text{N}$ *acinus-2 pinin-1*) | 5375 | 2936 | 94% | 5516 | 1.81 |
| Reverse labeling (R1)<br>($^{14}\text{N}$ *acinus-2 pinin-1*/$^{15}\text{N}$ Col) | 5415 | 3696 | 97% | 5740 | 1.60 |
| Reverse labeling (R2)<br>($^{14}\text{N}$ *acinus-2 pinin-1*/$^{15}\text{N}$ Col) | 5583 | 3745 | 97% | 5858 | 1.54 |
| Forward labeling (F1)<br>($^{14}\text{N}$ Col/$^{15}\text{N}$ *sec-5*) | 4655 | 4517 | 98.5% | 5361 | 0.96 |
| Forward labeling (F2)<br>($^{14}\text{N}$ Col/$^{15}\text{N}$ *sec-5*) | 5438 | 4855 | 98.5% | 6025 | 1.19 |
| Reverse labeling (R1)<br>($^{14}\text{N}$ *sec-5*/$^{15}\text{N}$ Col) | 5800 | 5922 | 98.5% | 6715 | 0.70 |
| Reverse labeling (R2)<br>($^{14}\text{N}$ *sec-5*/$^{15}\text{N}$ Col) | 5700 | 5977 | 98.5% | 6751 | 0.70 |

High-level labeling depends on three factors: 1) ^15^N containing salt needs to be over 99% purity; we find ^15^N chemicals from Cambridge Isotope Laboratories are generally high-purity. 2) The labeling time. We recommend growing *Arabidopsis* for 14 days to achieve high labeling efficiency. If plants can only be labeled for a shorter time before harvesting, then it is recommended to label the plants for one generation using a hydroponic system and start the experiment using the labeled seeds. If the Arabidopsis plants are small after 14 days of growth, then the labeling efficiency will be lower, for instance, our *acinus-2 pinin* mutants are smaller than wild-type plants, therefore the labeling efficiency is lower than wild-type with the same duration of labeling. 3) The availability of the ^15^N salt. Seeds should not be sown too many on solid-medium plates or in the liquid medium. We recommend the Arabidopsis plants are labeled 14 days or more to achieve high labeling efficiency and high identification rate, but users should be cautious not to stress plants by leaving them on medium for too long. Almost all proteins except seed storage proteins are labeled. These are not synthesized during the seedling stage and therefore they don’t incorporate the ^15^N labeling during growth and will remain unlabeled.

### 4.2 High-resolution scans for both MS^1^ and MS^2^ are key for high-quality data

Co-eluting peptides are common problems, especially in highly complex samples, and interfere with quantification. High-resolution scans in MS^1^ reduce peak overlap, improving the accuracy of quantification (Mann and Kelleher, 2008), so we typically acquire our data at 120K resolution.

High mass accuracy in MS^2^ helps to reduce the false discovery rate (FDR). Higher FDR was reported in the ^15^N sequence assignments due to more isobaric amino acid forms present in ^15^N labeling (Nelson et al., 2007) when the MS^2^ fragmentation was done using a low-resolution and mass accuracy QTOF2. To check this possibility in MS^2^ data acquired at high resolution, we compared the FDR in our seven labeled experiments listed in Table 2. After we combined peptides for ^14^N and ^15^N searches together with 1% FDR, we parsed the target and decoy ^14^N and ^15^N matches and calculated the FDR separately. We found the ^15^N data results had a lower FDR as that of ^14^N data search when MS^2^ scans were done in the high-resolution and high-mass accuracy Orbitrap mass spectrometer (Figure 4). This trend is more pronounced in the higher labeling efficiency datasets. If FDR was calculated based only on first three datasets (Col/*acinus-2 pinin-1*), there was no significant difference between ^14^N and ^15^N FDR, despite the average of FDR is slightly lower in ^15^N search. When four more Col/*sec-5* datasets were included for comparison, the ^15^N FDR is significantly lower than ^14^N one, indicating using high-resolution and high-accuracy measurement, the unique mass of ^15^N modification to the amino acid may empower less random matches in data searches.

**Figure 4.**
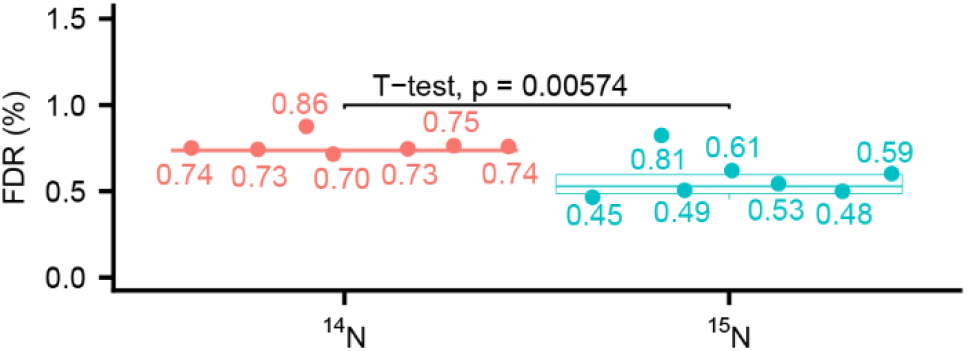
Comparison of false discovery rate (FDR) at peptide level between ^14^N search and ^15^N search shows ^15^N has a lower FDR. ^14^N and ^15^N searches are reported together allowing 1% FDR for peptides, then the target and decoy matches are parsed to calculate the FDR for ^14^N search and ^15^N search. Seven datasets (n=7) are used for the calculation.

### 4.3 Protein quantification process and result

After each peptide is quantified, they are compiled into protein groups in “Search Compare”. The spread of ratios for peptides from the same proteins are measured using the interquartile range, and Q1 (the lower quartile), median, and Q3 (the upper quartile) are reported, as illustrated in Figure 5A and 5B and quantification of the pairs can be visualized as Figure 5C and 5D). Here we include two biological experiments (including one forward and one reverse label of Col/*acinus-2 pinin-1* datasets) as a demonstration.

**Figure 5:**
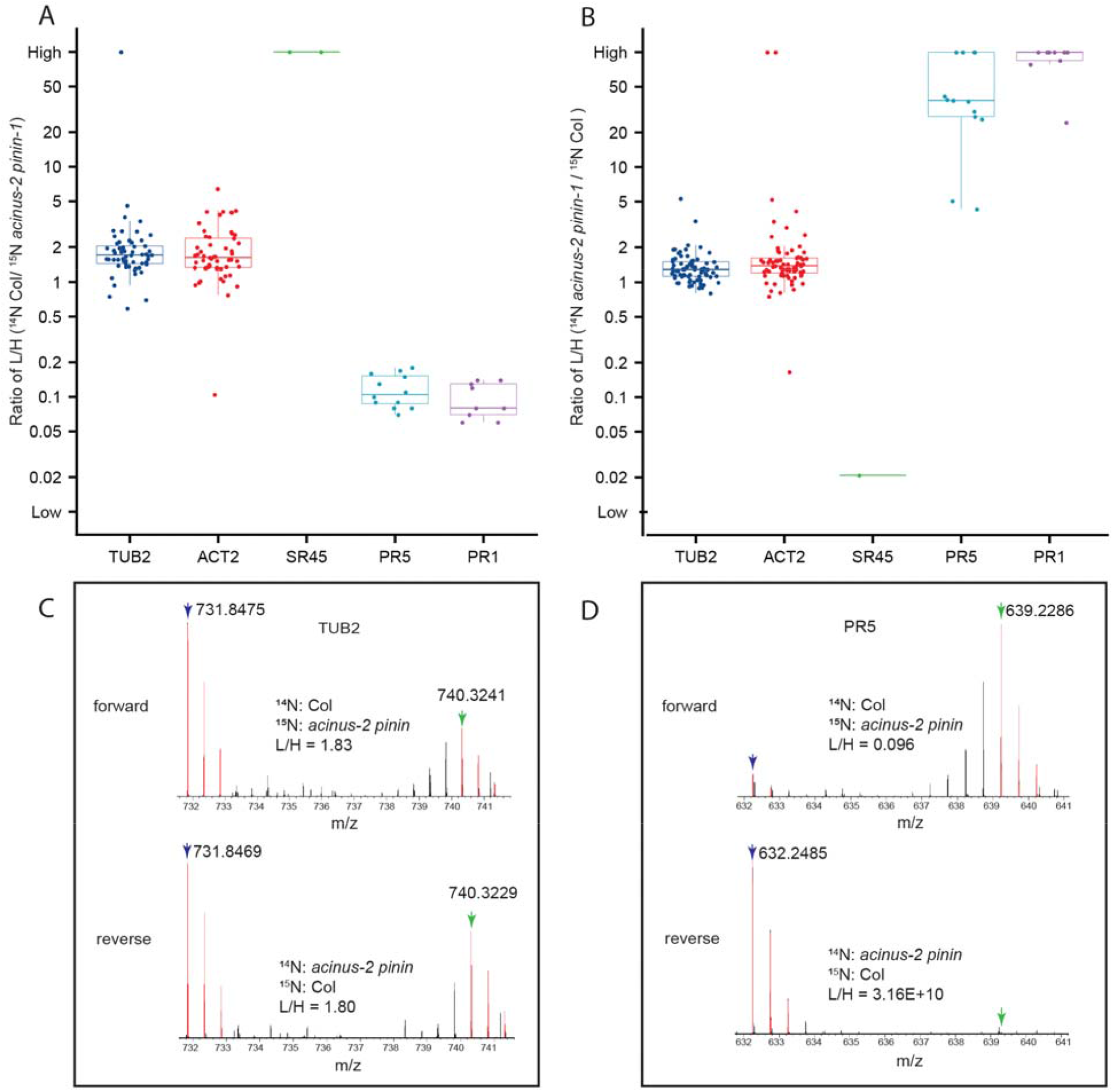
Quantitative ratio of proteins in wild-type Col and *acinus-2 pinin-1* double mutant before normalization. Each box plot shows the raw L/H labeled peptide ratios belonging to their appropriate protein as reported by Protein Prospector. (A) and (B) boxplot with beeswarm shown in two reciprocal labeled samples, SR45 is dramatically decreased, while PATHOGENESIS-RELATED GENE 5 and 1 (PR5 and PR1) show increased protein level in *acinus-2 pinin* double mutant. TUBULIN 2 (TUB 2) and ACTIN 2 (ACT2) are used as control. Three data points (a lower quartile (25%), median, and upper quartile (75%) are displayed in the output. Each dot is one peptide quantification. (C-D) MS1 spectra of the peptide “EVDEQMLNVQNK” from TUB2 and “FNTDQYCCR” from PR5 protein. Blue arrow: ^14^N labeled M peak; green arrow: ^15^N labeled M peptide. PR5 shows a significant increase in the double mutant. In the forward experiment, the ^15^N labeling efficiency is about 94%, while reverse labeling is about 97%.

To calculate the median and Q1, Q3, the log base 10 of all the ratios of the peptides from the same protein are first calculated in Protein Prospector to generate log base 10 of median and Q1, Q3, followed by converting these log values to normal values by raising 10 to the power.

To plot these quantification results, an R script is written as in Supplemental data 1. In the R script, the log base 2 of all the ratios of the peptides are calculated and converted back to normal values by ratio 2 to the power. The plot is the same as displayed in Protein Prospector, no matter base 10 or 2 is used.

A single protein can often be quantified by multiple peptides. A median value is preferentially reported instead of a mean value, as outliers, which are not unusual, can significantly skew the mean ratio, whereas median values are more tolerant. In general, the more peptides quantified from a single protein, the more accurate the median number is to the actual ratio. If the Q1 and Q3 are quite tight, then the quantification results are quite reliable. If the reciprocal labeling gives similar results, such as SR45, then the quantification should be reliable, even if the total peptides from this protein are only a few. We recommend at least three to four biological experiments be done for quantification, including at least one reciprocal labeling experiment (Wang et al., 2002).

### 4.4 Evaluation of quantification result

After the quantification is done, the users can evaluate the quality of the quantification of each protein and peptide of interest. Protein Prospector provides interactive feedback during the quantification process to allow for manual validation of the quantification results or visual assessment of what went wrong in case the ratio is incorrect. For protein quantification, a tight range between Q1 and Q3 often indicates the quantification is reliable. In cases where the range is big and the protein itself is of interest, then users can use the Cosine Similarity (CS) score to determine the quality of the matches or manually check them. CS scores are set up to determine the quality of the matching (L+H) peaks. Once the peptide sequence is identified, the elemental composition of the peptide is generated based on the peptide sequence. The CS score, similar to the Isotope Dot Product “idotp” product used in Skyline (Schilling et al., 2012), automatically measures the similarity between the experimentally measured isotope pattern and the calculated pattern using the M, M+1, M+2 peaks, thus reducing manual checking time by auto-flagging the contaminated peaks (Figure 6). The CS score ranges between 1.0 (a perfect match) and 0.0 and can be determined by measured peak intensity or area. Figure 6A shows the pair have both good CS scores and the L/H ratio of this peptide is close to the median number of the protein. Figure 6B shows one peptide in the pair of another peptide from the same protein gives a lower CS score (the M peak is contaminated with a co-eluting peptide), and thus the L/H ratio of this peptide will produce an outlier ratio.

**Figure 6:**
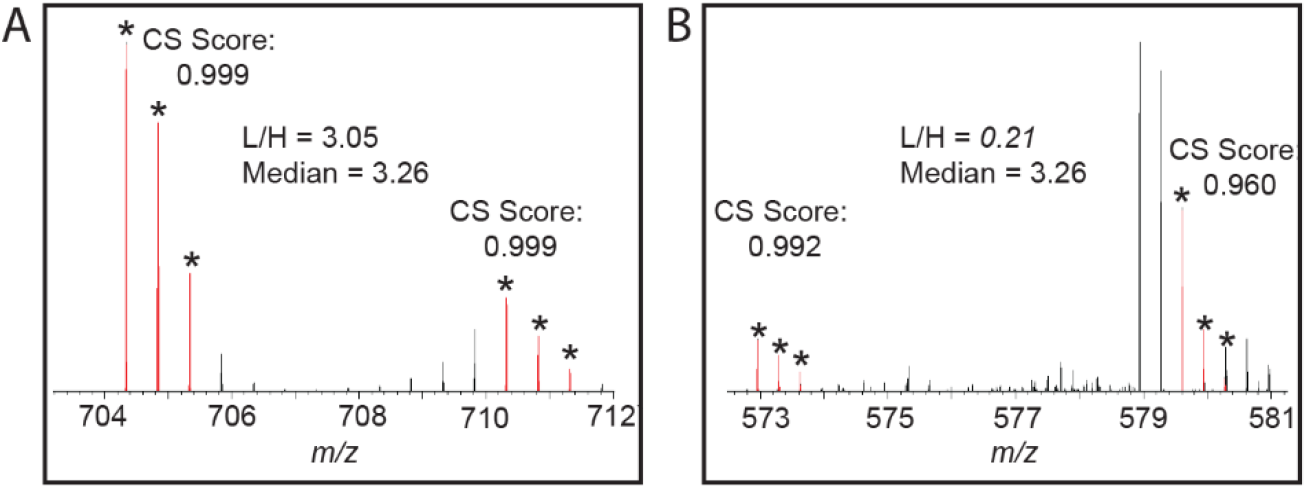
(A) Good CS scores are reported for the ^14^N and ^15^N peptide pair of “LTYYTPEYETK” from the protein RIBULOSE-BISPHOSPHATE CARBOXYLASE (RBCL) in MS^1^ and the L/H ratio of this peptide is close to the median ratio. (B) The L/H ratio of the peptide “DLAVEGNEIIREACK” from RBCL shows outlier quantification measurement and a poor CS score is reported for ^15^N labeled peptide in this pair.

Importantly, Protein Prospector takes account of labeling efficiency when calculating the CS score, as low labeling efficiency changes the isotope pattern quite dramatically. It should be noted that the CS score will be less accurate when the peak intensity of the peptide is very low.

### 4.5 Analysis data storage and quick data retrieved enabled by cache function

Users can create a cache file when submitting the quantification in Search Compare. This stores the data required to regenerate the Search Compare report in a JSON file. The cache function is quite useful for various reasons: 1) when the user needs to retrieve the data or manually check the data, there is no need to re-calculate the quantification, which can take many hours for a large dataset. With the cache file, the reports come up quickly for a few seconds rather than hours; 2) Often it is hard to display an HTML peptide report when many proteins or peptides are quantified. The cache function can allow visualization of such reports easily.

### 4.6 Applications and limitations, further steps and perspectives

This workflow allows users to report quantification of thousands of proteins and is applicable to the quantification of the total proteomes, sub-proteomes, and immunoprecipitated samples (Bi et al., 2021; Garcia et al., 2020; Park et al., 2019). During the extraction of elution profile of every peptide identified, Protein Prospector averages together scans over a time window but doesn’t fit the peak shape to a Gaussian function. Therefore, each identified peptide/protein will be quantified and none gets discarded due to failing the scoring threshold for fitting the Gaussian function.

To get high quality data, we recommend to get 97% or above labeling efficiency to achieve higher ID rates in ^15^N samples, so more proteins will be reproducibly identified and quantified between different replicates. Data acquired on high resolution and high accuracy instruments will also improve the quality of the dataset.

A systematic normalization is normally required before comparing results between different experiments (Ting et al., 2009), as the samples are rarely mixed at exactly 1:1. One choice is to use the median number of all the quantified proteins, or median number of top one hundred abundant proteins (Table 2). Alternatively, users can use housekeeping proteins that are assumed to not change for normalization.

Statistical analysis of quantification data on three or more replicates is advised. Users need to determine how to apply statistical analysis on the data using a separate tool. Benjamini-Hochberg (BH) multiple hypothesis test has been used to determine significant regulated PTM peptide groups in ^15^N metabolic labeled samples (Wong and Li, 2021). A standardized statistic pipeline for protein quantification is still lacking, particularly a pipeline that can leverage quantification ratios of each peptide from a protein. Our current workflow uses a median value, which takes advantage of the quantification ratios of each peptide but is less affected by outliers than using the mean. However, the statistical power utilizing quantifications from these multiple peptides from single protein has not been explored and awaits development in the future.

A targeted quantification strategy is recommended for further analysis of proteins of interest because this provides more accurate quantification and is less likely to have missing values, particularly in the ^15^N labeled samples (Bi et al., 2021; Reyes et al., 2021). In addition to targeted analysis, data-independent acquisition (DIA) (Zhang and Bensaddek, 2021) can also be utilized, which can be done in label free samples or combined with ^15^N metabolic labeling in the future. DIA benefits from having few missing values, but more efforts will be needed to deconvolve the mixed MS^2^ spectra in DIA datasets.

^15^N metabolic labeling has been utilized in studies of analyzing protein synthesis and degradation (Guan et al., 2011a; Guan et al., 2011b). These studies are based on incomplete and often low incorporation rates which result in very broad satellite peak distributions and cause ^15^N labeled peptide isotope clusters to overlap with ^14^N labeled peptide clusters. As Protein Prospector doesn’t deconvolve the ^15^N distribution from the ^14^N distributions (separate the ^14^N and ^15^N signal when peaks overlap), the presented workflow will not provide accurate quantification in this type of study. On the other hand, for chase studies that analyze the assembly kinetics *in vitro* (Bunner and Williamson, 2009), the presented workflow can be applied because the proteins involved have a very high labeling efficiency.

This workflow can be also applied to the quantification of post-translational modification with a slight modification. The users will include related PTM search parameters into data search. Instead of reporting median number at protein level, ratios from each peptide are reported and then compared across different replicates.

## Data Availability Statement

The mass spectrometry proteomics data for this study have been deposited to the ProteomeXchange Consortium via the PRIDE partner repository. Col/*acinus-2 pinin-1* and Col/*sec-5* data are available via ProteomeXchange with identifier PXD030081 and PXD030575.

## Author Contributions

R.S., S.-L.X. designed the experiment and R.S. performed the experiment, R.S., A.V.R., and S.-L.X. analyzed the data sets and generated figures. P.R.B. and R.J.C. provided technical support for Protein Prospector and revised the manuscript. Z.Y.W. provided suggestions. R.S., A.V.R. and S.-L.X. wrote the manuscript.

## Funding

This research was funded by the NIH grant R01GM135706 to S.-L.X. and Carnegie endowment fund to Carnegie mass spectrometry facility.

## Conflict of interest

The authors declare that the research was conducted in the absence of any commercial or financial relationships that could be construed as a potential conflict of interest.

## Acknowledgments

We thank Drs. Ajeet Chaudhary and Sumudu Karunadasa for critical reading of the manuscript.

## Supplemental Material

The Supplemental figure 1, 2 and Supplemental data 1 are provided.

---

**Supplemental Figure 1:**
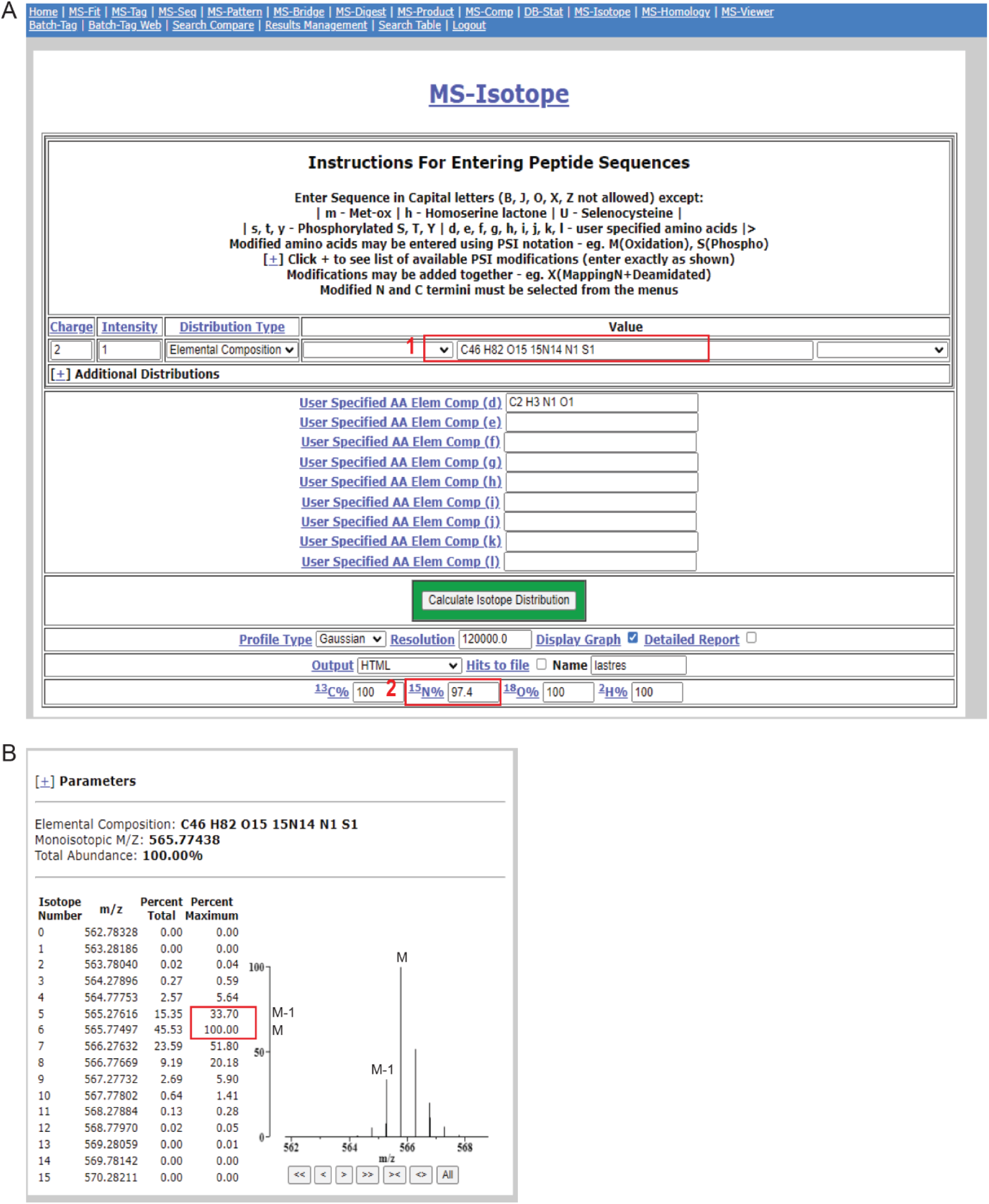
Using MS-Isotope module in Protein Prospector to generate theoretical ^15^N peak pattern. **(A)** Users need to input either composition [1] or sequence of the peptide (not shown) and input the 15N incorporation rate [2]; (B) Protein Prospector will generate theoretical peaks and percentage number based on input from A.

**Supplemental Figure 2:**
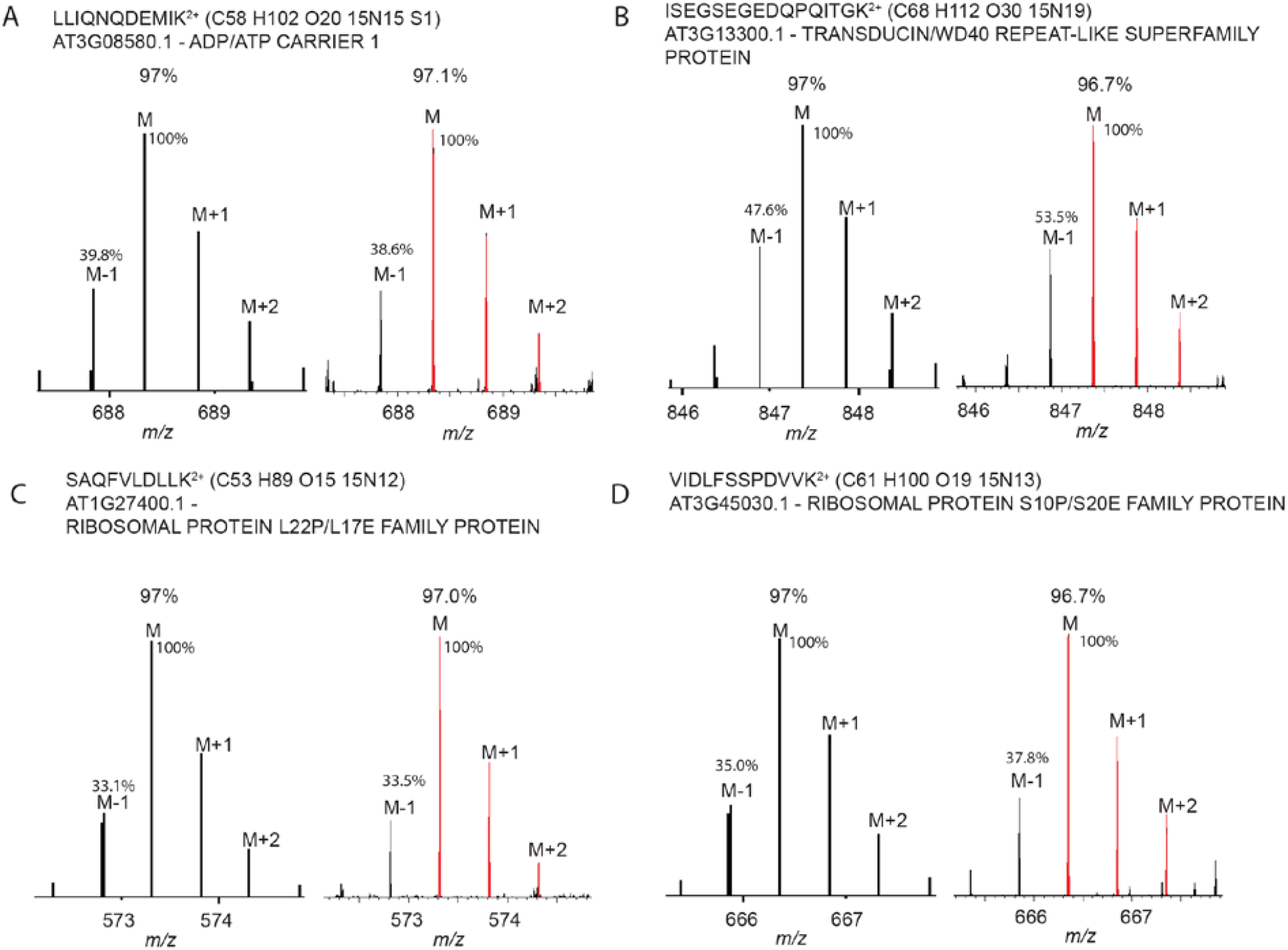
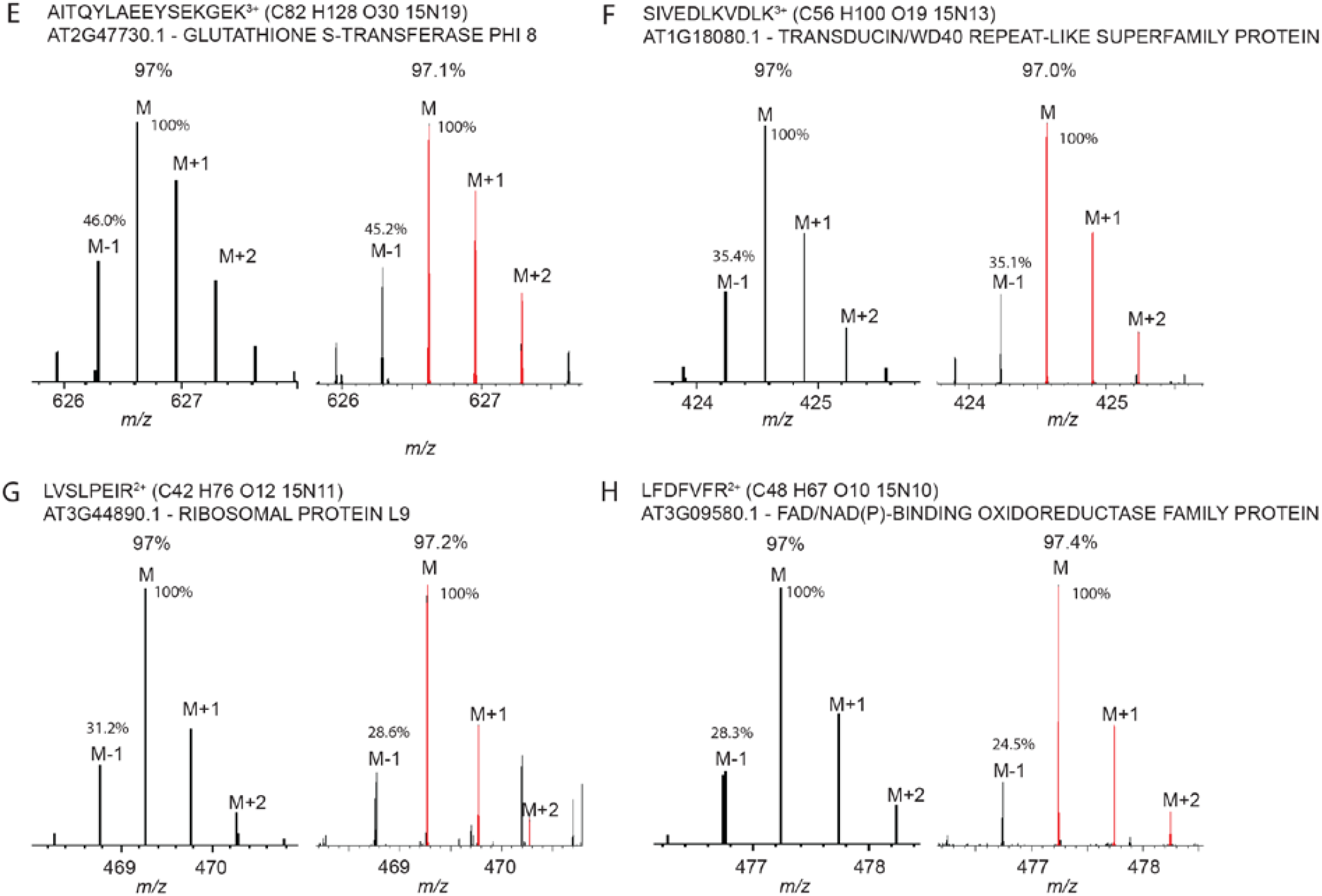
Incorporation of the ^15^N into the Arabidopsis Col samples was 97%. (A-H) Eight ^15^N peptides from different proteins were examined to determine the labeling efficiency, with the left showing theoretical peak pattern at 97% labeling efficiency for all the peptides, and the right (M, M+1, M+2, colored in red) showing the experimental peak pattern and calculated labeling efficiency. Smaller *m/z* peptides are more preferred to determine the labeling efficiency, because the monoisotopic peak (M) is the largest peak and can be set as 100% maximum during calculation.

**Supplemental Data 1:**
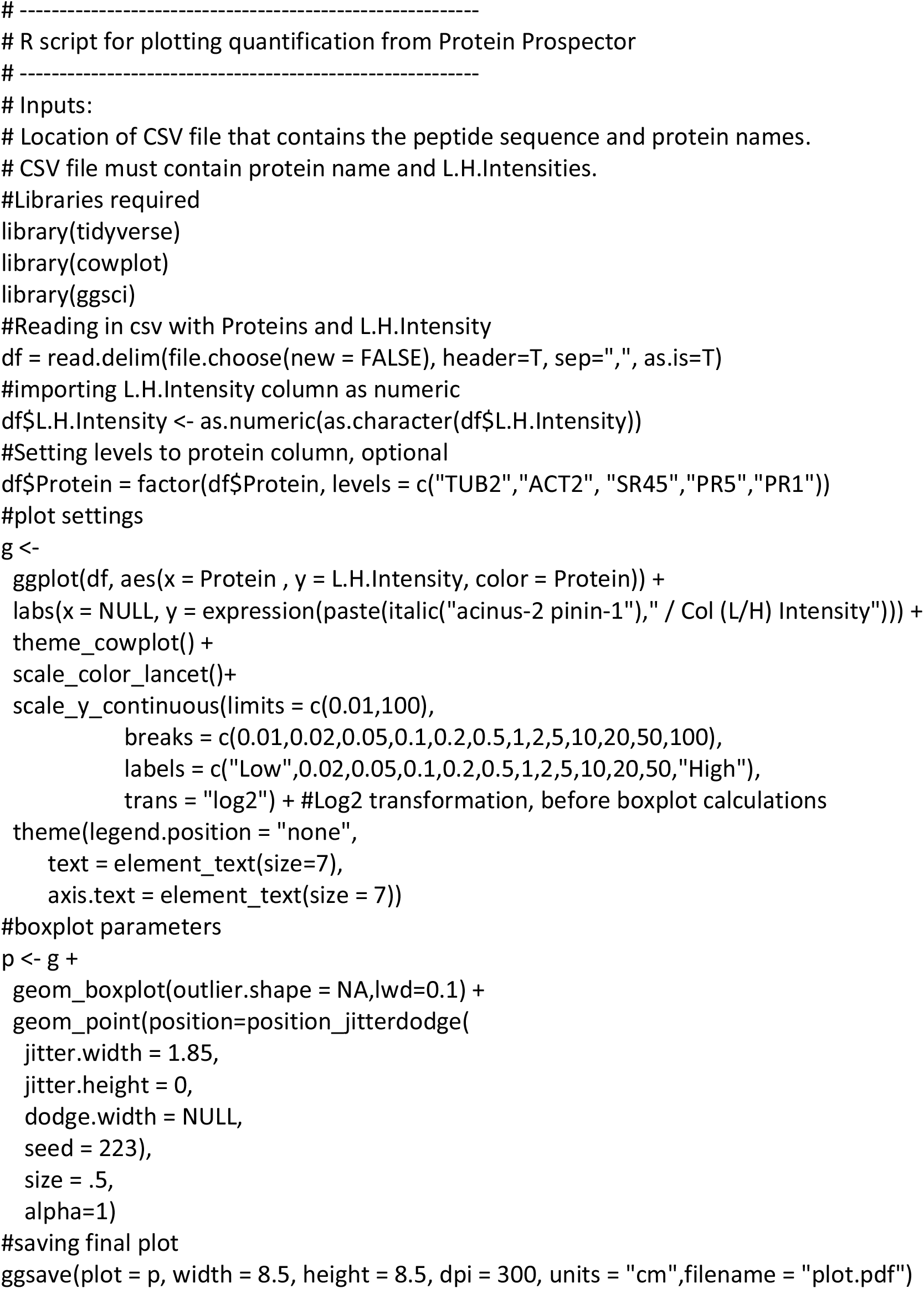
R script used to generate the median and Q1/Q3 for quantification.

## Notes

### Competing Interest Statement

The authors have declared no competing interest.

